# Single Cell Profiling in the *Sox10^Dom/+^* Hirschsprung Mouse Implicates *Hoxa6* in Enteric Neuron Lineage Allocation

**DOI:** 10.1101/2024.09.18.613729

**Authors:** Justin A. Avila, Joseph T. Benthal, Jenny C. Schafer, E. Michelle Southard-Smith

## Abstract

**Background & Aims:** Enteric nervous system (ENS) development requires migration, proliferation, and appropriate neuronal diversification from progenitors to enable normal gastrointestinal (GI) motility. *Sox10* deficit causes aganglionosis, modeling Hirschsprung disease, and disrupts ratios of postnatal enteric neurons in proximal ganglionated bowel. How *Sox10* deficiency alters ratios of enteric neuron subtypes is unclear. *Sox10’s* prominent expression in enteric neural crest-derived progenitors (ENCP) and lack of this gene in enteric neurons led us to examine *Sox10^Dom^* effects ENS progenitors and early differentiating enteric neurons.

**Methods:** ENS progenitors, developing neurons, and enteric glia were isolated from *Sox10^+/+^* and *Sox10^Dom/+^* littermates for single-cell RNA sequencing (scRNA-seq). scRNA-seq data was processed to identify cell type-specific markers, differentially expressed genes, cell fate trajectories, and gene regulatory network activity between genotypes. Hybridization chain reaction (HCR) validated expression changes detected in scRNA-seq.

**Results:** scRNA-seq profiles revealed three neuronal lineages emerging from cycling progenitors via two transition pathways accompanied by elevated activity of *Hox* gene regulatory networks (GRN) as progenitors transition to neuronal fates. *Sox10^Dom/+^* scRNA-seq profiles exhibited a novel progenitor cluster, decreased abundance of cells in transitional states, and shifts in cell distributions between two neuronal trajectories. *Hoxa6* was differentially expressed in the neuronal lineages impacted in *Sox10^Dom/+^*mutants and HCR identified altered *Hoxa6* expression in early developing neurons of *Sox10^Dom/+^* ENS.

**Conclusions:** *Sox10^Dom/+^* mutation shifts enteric neuron types by altering neuronal trajectories during early ENS lineage segregation. Multiple neurogenic transcription factors are reduced in *Sox10^Dom/+^* scRNA-seq profiles including multiple *Hox* genes. This is the first report that implicates *Hox* genes in lineage diversification of enteric neurons.

## Introduction

The Enteric Nervous System (ENS) is a complex network composed of the myenteric and submucosal ganglia within the intestinal wall. This interconnected network is essential for gastrointestinal (GI) function, including regulation of bowel motility and maintaining intestinal homeostasis^1^. Normal ENS function relies on the complex interplay of more than 20 distinct neuron types comprised of excitatory, inhibitory, interneuron, and sensory neuron classes^2-4^. The gene regulatory networks (GRN) that direct the development and differentiation of these neuronal subtypes are not well understood.

Initial development of the ENS relies on temporally and spatially regulated migration and differentiation of enteric neural crest-derived progenitors (ENCP)^5-7^. After separating from the neural tube, vagal ENCPs migrate into the developing fetal intestine, initiating rostral-to-caudal colonization of the gut that is complete by 14.5 days post-coitus (dpc) in the mouse. Concurrent with ENCP colonization, differentiation and maturation of enteric neurons and glia occurs and continues through postnatal maturation of the ENS^8^. Disruption of these complex processes can produce GI motility disorders and aberrant bowel function.

The necessity of normal ENS function and the consequences of abnormal enteric neurons are evident in patients with Hirschsprung’s disease (HSCR). HSCR is a congenital disorder diagnosed by absence of enteric ganglia in a variable portion of the distal bowel due to incomplete ENCP colonization of the developing gut. The lack of nerve cells in the HSCR aganglionic segment predisposes to life-threatening intestinal blockage. Current therapies for HSCR rely on surgical removal of aganglionic regions. Despite removal of aganglionic segments, many HSCR patients suffer residual GI motility issues. Realization that altered ratios of enteric neuron types in proximal ganglionated intestine of HSCR mice and patients underlies continued aberrant bowel motility has emphasized the need for understanding regulatory mechanisms that control enteric neuron lineage diversification^9, 10^.

*Sox10* is an essential transcription factor for ENS development that is crucial for initial colonization of the gut by vagal ENCPs and plays distinct roles in developing neuronal and glial populations within the ENS^11^. Deficiency of *Sox10* causes delays of vagal ENCP migration along the developing fetal intestine^12^, leading to aganglionosis of the distal bowel, the defining feature of Hirschsprung disease. In prior work we documented the effect of *Sox10* on the mature enteric neuron subtypes in the postnatal ENS^9^. Ratios of calretinin-expressing motor neurons within the duodenum and ileum of *Dominant megacolon* mice (*Sox10^Dom/+^*) were significantly altered, revealing *Sox10’s* effect on allocation of enteric neuron ratios in the mature ENS^9^. Because the expression of *Sox10* is restricted to developing ENCPs and later to enteric glia in the postnatal ENS, these findings raise questions of how *Sox10* deficit alters neuronal distributions since the gene is not expressed in neurons. Known roles for *Sox10* in regulating differentiation of trunk NC-derived cells and maintaining the neurogenic and gliogenic potential of cultured ENCPs *in vitro^13, 14^* suggest early actions of this transcription factor may feed forward into later processes of cell fate diversification.

Knowledge of cellular diversity and relationships between progenitors and emerging neurons in ENS development has been greatly enhanced by single-cell RNA sequencing (scRNA-seq) technologies^15, 16^. Transcriptional profiling has consistently identified two major populations of cells in the developing ENS at 15.5 dpc: cycling progenitor cells and differentiating lineages. Cycling ENCPs are comprised of undifferentiated cells expressing known ENS progenitor markers. The second major population detected at 15.5 dpc consists of differentiating cells transitioning toward neuronal fates with smaller numbers of emerging enteric glia. Among the neuronal precursors, two primary trajectories or “branches” have been reported that are characterized by the expression of genes including NO synthetase (*Nos1*), Vasoactive intestinal peptide (*Vip*), or Choline acetyltransferase (*Chat*) ^4, 17^. Analysis of these scRNA-seq data have identified regulatory roles for *Pbx3* and *Sox6* for progression of ENCPs into enteric neurons^4, 18^. These studies illustrate the value of scRNA-seq for mapping developmental trajectories and revealing genes associated with cell type specification in the developing ENS.

To better understand neuronal diversification in the ENS, we employed scRNA-seq analysis to examine cellular dynamics and gene regulatory networks in the *Sox10^Dom/+^* HSCR mouse model. We hypothesized that dysregulation of *Sox10* in ENCPs would disrupt gene regulatory networks required for enteric neuron lineage diversification. By combining two transgenic reporter lines with the *Sox10^Dom/+^* model, we successfully isolated ENCPs and differentiating neuronal and glial lineages at 15.5 dpc for single cell transcriptomics. Analysis of the resulting scRNA-seq data unexpectedly identified three distinct neuronal trajectories, indicating greater complexity during enteric neurogenesis than previously appreciated. In *Sox10^Dom/+^* mutants, cell distributions are altered between two of the three developing neuronal trajectories revealing a significant role for *Sox10* in governing early enteric neuron lineage allocation. A unique progenitor cluster was also identified in *Sox10^Dom/+^* populations accompanied by a reduction in calculated transition rates as progenitors progress towards neuronal fates. Differential gene expression analysis between *Sox10^+/+^* and *Sox10^Dom/+^*data identified downregulation of multiple positive transcriptional regulators associated with neurogenesis. Gene regulatory network analysis indicated heightened Hox-associated regulon activity within neuronal lineages. Moreover, we found *Hoxa6* transcription was specifically disrupted in *Sox10^Dom/+^* scRNA-seq profiles and alterations in *Hoxa6* mRNA *in situ* were confirmed by Hybridization Chain Reaction (HCR). These findings highlight the importance of *Sox10* expression during early neuronal specification and contribute to the broader understanding of cellular diversification during ENS development.

## Results

### Multiple ENS Lineages Are Captured Via a Dual Transgenic Approach Utilizing *Sox10*-YFP and *Phox2b*-CFP Reporters

Prior labeling of enteric ganglia with the canonical *Wnt1*-cre transgene did not comprehensively mark all *Phox2b*+ neurons^19^. In contrast, a *Phox2b*-CFP transgene was observed to label all HuC/D+ enteric neurons^20^. Therefore, we utilized a combinatorial transgenic approach to capture the full profile of ENCPs, differentiating enteric neurons and glia that relied upon the transgenic lines *Phox2b*-CFP and *Sox10-*YFP^20, 21^. This strategy enabled simultaneous capture of ENCPs (YFP+ / CPF+) as well as emerging enteric glia expressing low intensity CFP (YFP+, CPF^Low^+) and differentiating enteric neurons exhibiting high CFP intensity (CPF^High^+) from the *Phox2b*-CFP transgene (Figure 1A). Fetal intestines from crosses with *Sox10^Dom/+^* mutants were collected at 15.5dpc and examined for fluorescent expression to score the extent of bowel colonization. Since *Sox10^Dom/+^* aganglionosis can be highly variable^22^, we selected samples exhibiting total colonic aganglionosis for tissue dissociation (Figure 1B) to increase the likelihood of detecting effects due to the *Sox10^Dom/+^* allele. Subsequently, dissociated cell suspensions were flow sorted to isolate viable cells that were expressing either the CFP or YFP transgene from four *Sox10^+/+^* fetal intestines and three *Sox10^Dom/+^* samples for scRNA-seq.

**Figure 1:**
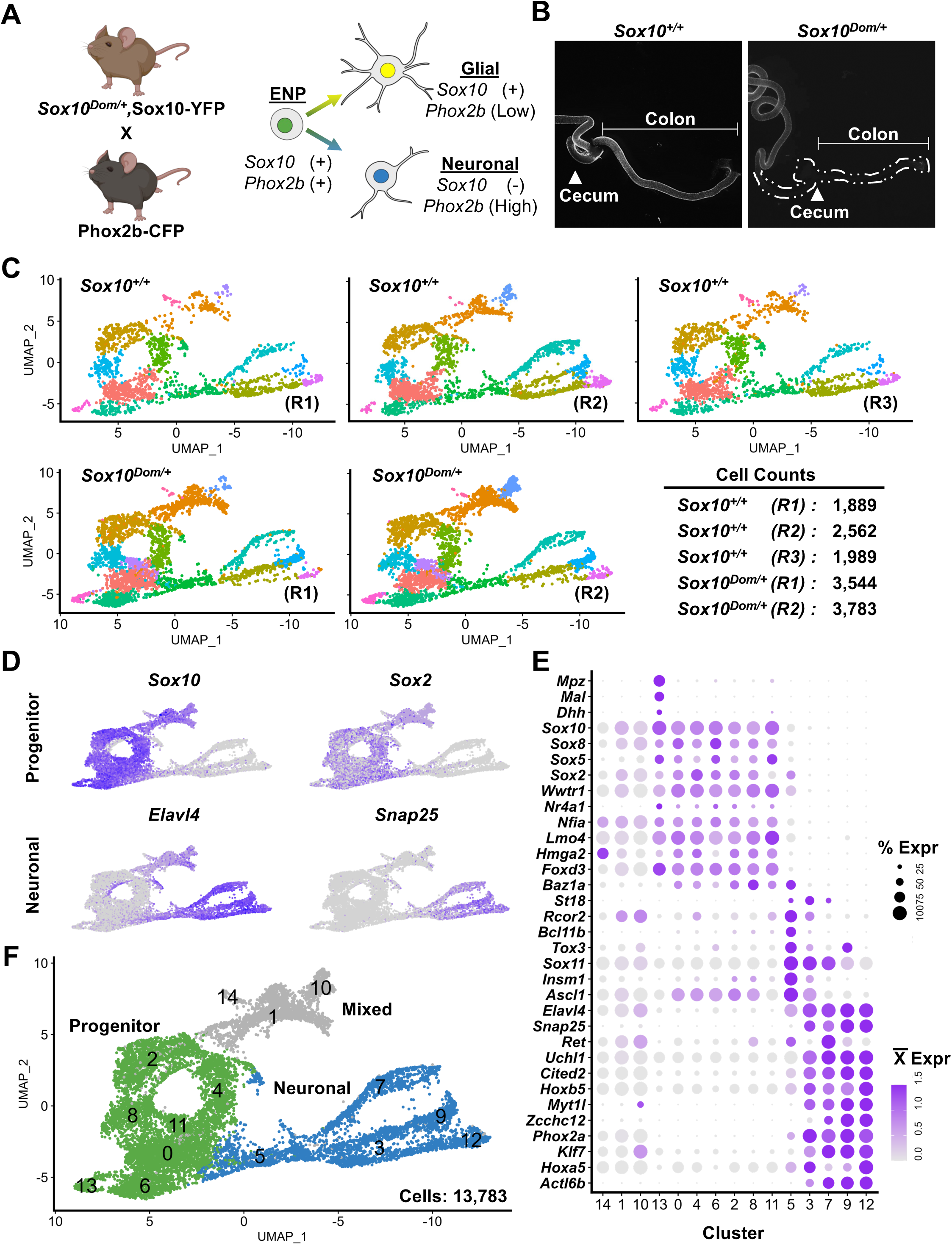
Dual *Sox10-Phox2b* transgene strategy comprehensively captures developing ENS populations. A) Transgenic lines crossed to enable flow sort isolation of ENS lineages marked by expression of *Sox10*-YFP and *Phox2b*-CFP in progenitors (YFP+/CFP+), maturing neurons (CFP^high^), and enteric glia (YFP+, CFP^low^). B) Wholemount fluorescent images of YFP expression in *Sox10^+/+^* and *Sox10^Dom/+^* fetal intestines at 15.5 dpc. Dotted lines emphasize the hindgut borders in *Sox10^Dom/+^* mutant where YFP cells are lacking. C) UMAP representations display scRNA-seq data generated for biological replicates of *Sox10^+/+^* and *Sox10^Dom/+^* accompanied by cell counts for each replicate. D) UMAP representations of marker genes for progenitors (*Sox10*, *Sox2*) and developing neurons (*Elavl4*, *Snap25*). E) Dot plot of known marker genes for ENS progenitors, neuroblasts, and neurons per cluster. F) UMAP of clusters shared by *Sox10^+/+^* and *Sox10^Dom/+^*classified by marker gene expression and Gene Ontology analysis identified three main groups: progenitors (green), neurons (blue), and mixed (gray).

To assess similarities and differences between biological replicates of *Sox10^+/+^* and *Sox10^Dom/+^* single cell populations, we compared Uniform Manifold Approximation and Projections (UMAPs) between genotypes after integration, merging and data normalization^23^. Based on run quality and appropriate integration three of the four *Sox10^+/+^* biological replicates (6440 cells) and two of the three *Sox10^Dom/+^* data sets (7,327 cells) appeared similar and were analyzed further (Figure 1C).

Expression patterns of known cell type marker genes were examined in the merged *Sox10^+/+^* and *Sox10^Dom/+^* datasets to ascertain cell identities. Feature plots of progenitor genes (*Sox10*, *Sox2*) and neuronal genes (*Elavl4*, *Snap25*) were used initially to demarcate the UMAP spatial distribution of progenitors and differentiating neurons (Figure 1D). Extended analysis of established marker genes confirmed the identity of these populations as progenitors (clusters 0, 2, 4, 6, 8, 11, 13) and neurons (clusters 3, 5, 7, 9, 12) (Figure 1E and 1F). Gene ontology terms based on differentially expressed “marker” genes for these clusters were also consistent with progenitor and neuronal identities (Tables 1 and 2). Cluster 13 exclusively expressed myelination markers *Mpz* and *Mal* and elevated expression of *Dhh*, identifying these cells as nerve-associated neural crest progenitors^24^ (Figure 1E). Clusters 1, 10, and 14 exhibited mixed expression of both neuronal and glial genes (Figure 1E) concurrent with high 18s rRNA gene expression, high mitochondrial and low RNA feature counts that led us to exclude these clusters from further analysis. The number of high-quality cells and the presence of appropriate cell identities confirm the successful isolation of developing ENS lineages in wildtype and *Sox10^Dom/+^* littermates.

### scRNA-seq Profiling at 15.5 dpc Identifies Unexpected Complexity of Early ENS Neuronal Lineages

In total, 5,583 *Sox10^+/+^* cells and 5,549 *Sox10^Dom/+^* cells, comprising progenitor and neuronal populations, were subset totaling 11,132 cells. We performed unsupervised sub-clustering to assess potential for additional cell states. Overall, nineteen sub-clusters were evident in the combined *Sox10^+/+^* and *Sox10^Dom/+^* data that were classified as neuronal (NC) or progenitor (PC) (Figure 2A). Neuronal cells resolved into twelve sub-clusters that made up three separate neuronal branches. To date, prior scRNA-seq studies of the ENS have identified two neuronal lineages at 15.5 dpc^4^. To evaluate whether the third neuronal branch was inherent to the data or could possibly be attributed to the analysis approach, we compared our datasets with other ENS scRNA-seq data^4^. Comparable reprocessing of Morarach’s 15.5 dpc dataset identified distinct progenitors and neuronal populations with two separate neuronal branches as previously reported. We then integrated our *Sox10^+/+^* with the reprocessed Morarach 15.5 dpc dataset to assess whether the integration of these datasets led to consolidation of the three *Sox10^+/+^* neuronal lineages into two. Noticeable similarity was apparent in initial PCA plot distributions between both datasets. After Harmony batch correction to account for technical variations (ex., experimental conditions, sample processing, or sequencing procedures)^25^, improved alignment of principal component embedding values was seen indicating integration (Figure 2B). The Harmony UMAP analysis revealed that cell distributions, consisting of progenitor and neuronal lineages, are evenly distributed except for a subset of *Sox10^+/+^* cells (Figure 2C, arrow). This third branch is still separated upon integration with the Morarach data and does not collapse back into two upon integration. Thus, our data indicate the existence of a third neuronal lineage at 15.5 dpc (Figure 2D). Moreover, the presence of this third branch in the *Sox10^+/+^* data suggests the third trajectory is a consequence of the dual transgene strategy because the Morarach data did not separate into three branches upon integration.

**Figure 2:**
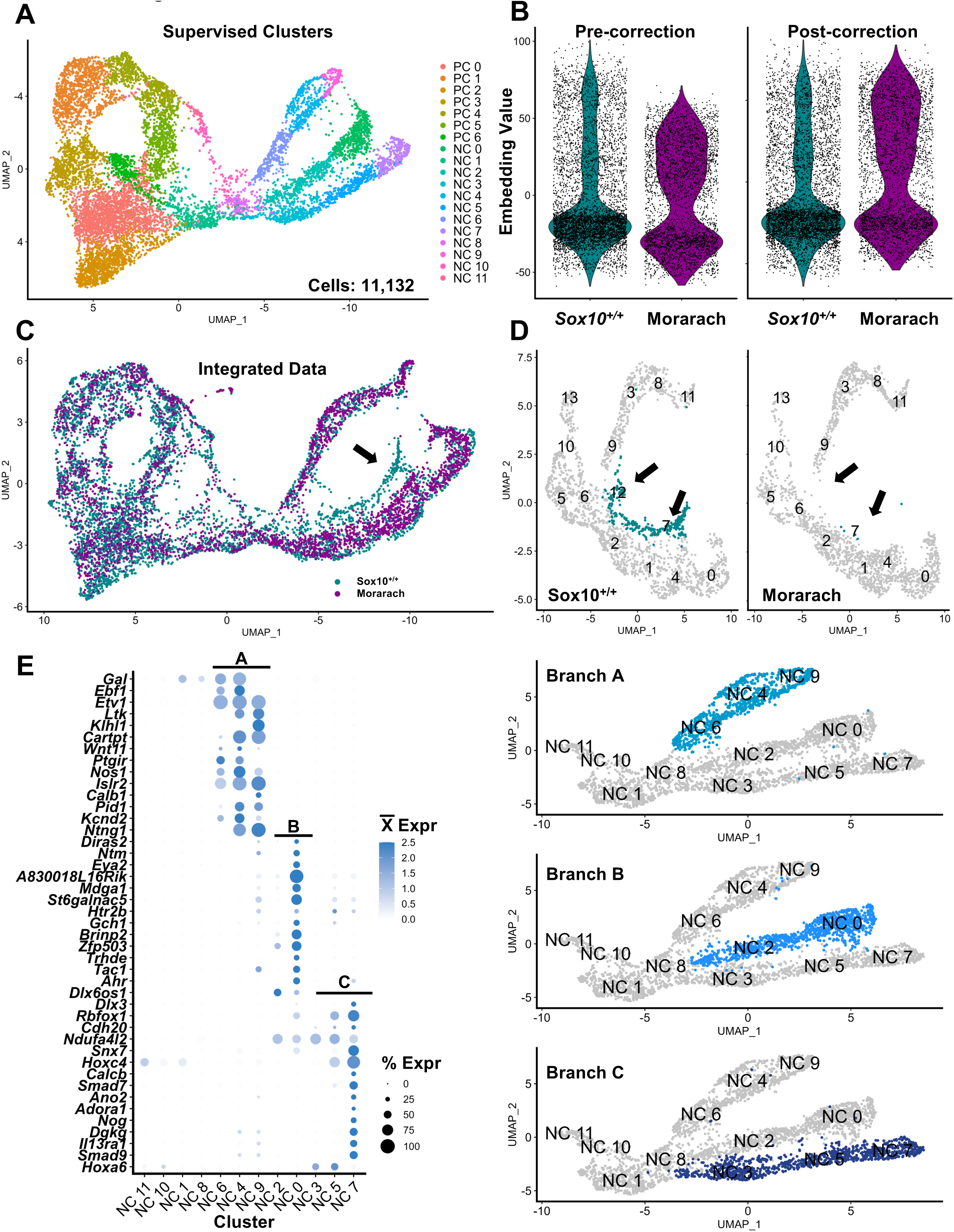
scRNA-seq detects earlier diversity of neuronal lineages than previously appreciated. A) UMAP visualization displays 19 distinct clusters detected during supervised clustering of progenitor and neuronal populations. These clusters were subsequently tracked as progenitor clusters (PC 0-6) and neuronal clusters (NC 0-11). B) Violin plots show pre- and post-Harmony batch correction for the embedding values of this study’s *Sox10^+/+^*data and the Morarach et al., 15.5 dpc dataset. C) UMAP showing the dataset overlay after Harmony batch correction color-coded by dataset origin. D) UMAPs display the subset and reclustered analysis of neuronal populations from batch-corrected data. Arrows are used to indicate differences in cluster 12 and 7 distributions highlighted in teal. E) Dot plot displays gene expression detected in the neuronal lineages for the aggregated *Sox10^+/+^* and *Sox10^Dom/+^* data. Minimum average expression set to 0. Adjacent UMAPs highlight the specific branches designated as Branch A (NC6, NC4, and NC9), Branch B (NC2 and NC0), and Branch C (NC3, NC5, and NC7).

### Branch-Restricted Marker Genes in Early Enteric Neuron Lineages Are Expressed in Later Development and Maintained in Mature Enteric Neurons

One of the challenges in enteric neurobiology has been tracing discrete neuron types from their first emergence to adulthood. Elegant studies have traced the initial cell cycle exit of enteric neurons based on neurotransmitters or birth dating with nucleotide analogs^26-28^. However, there are some inconsistencies between the early appearance of neuron markers like Nitric Oxide Synthase (NOS1) and the final phenotype of some enteric neuron types. To evaluate genes expressed in early differentiating enteric neuronal lineages as potential markers for labeling subsequent mature enteric neurons, we examined whether marker genes present within each neuronal branch at 15.5 dpc are expressed over the course of ENS development and maturation. Potential marker genes for each branch were first identified using the FindAllMarkers function in Seurat for each neuronal cluster in the combined *Sox10^+/+^* and *Sox10^Dom/+^*data. We then manually examined the expression levels and spatial distribution of the upregulated genes within each cluster to prioritize those that distinguish the three neuronal branches A (NC6, NC4, and NC9), B (NC2 and NC0), and C (NC3, NC5, and NC7) (Figure 2E, Table 3). Branch-specific marker genes limited to the termini of neuronal branches A (NC9), B (NC0), and C (NC7) at 15.5 dpc were prioritized. We subsequently evaluated whether these branch-specific markers were detected in later fetal stages or adult enteric neurons by screening for expression in other publicly available ENS datasets^3, 4^. We observed that the co-expression of terminal Branch A markers (*Islr2* and *Cartpt)* is limited to distal branch clusters and *Cartpt* is maintained in subsets of mature neurons. Similar restricted expression is also observed for terminal cluster genes in Branch B (*Gch1* and *Tac1*) and Branch C (*Calcb* and *Nog*) (Figure 3). These branch-restricted marker genes may be useful for future developmental tracking of distinct neuron types.

**Figure 3:**
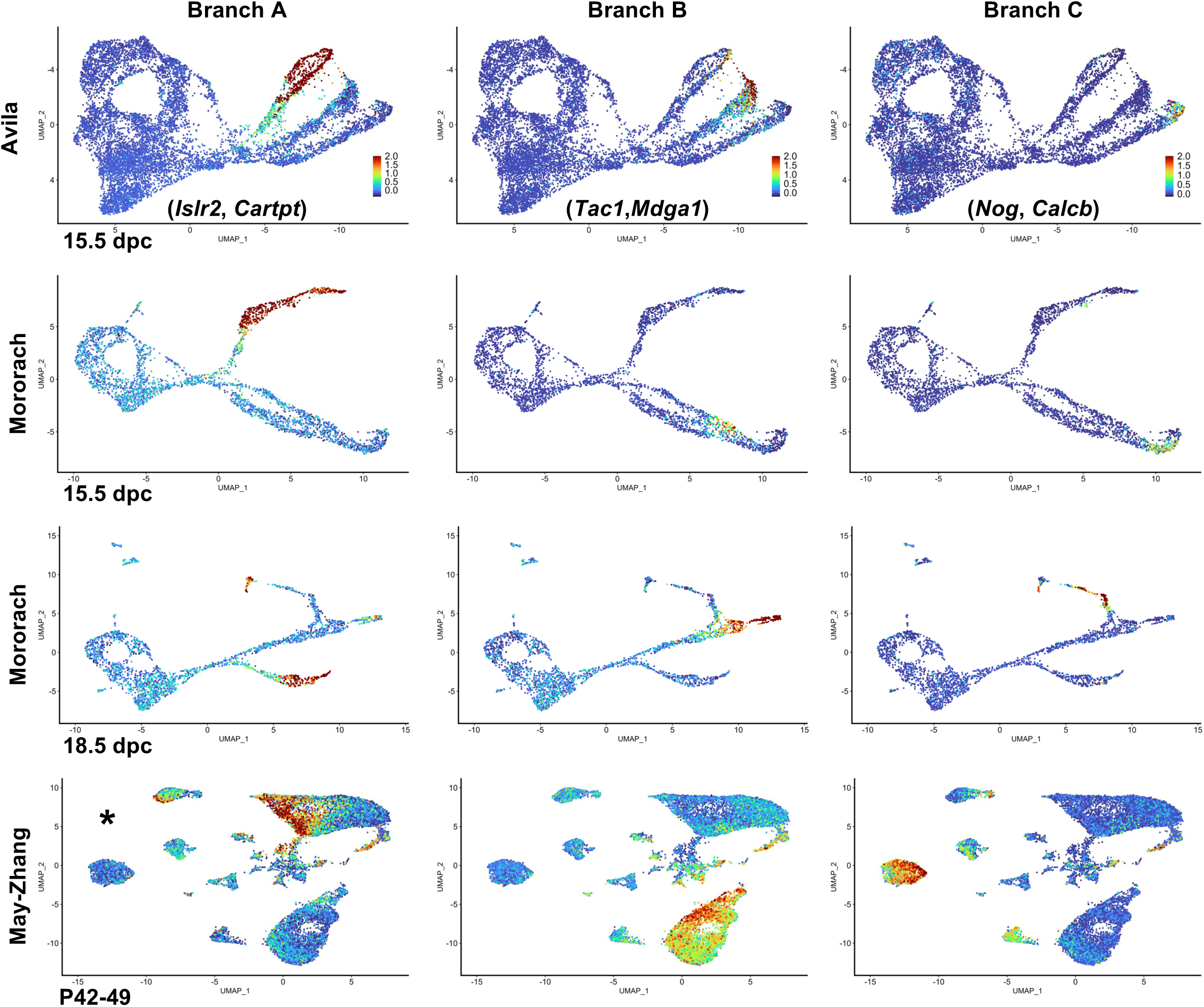
Neuronal-specific markers persist and highlight distinct cell populations throughout ENS development. Feature plots demonstrating pairs of co-expressed markers restricted to each neuronal branch for multiple enteric single-cell data sets across later fetal stages (Morarach et al., data at 15.5dpc and 18.5dpc)^4^ and into adulthood (May-Zhang et al., data at P42-49)^3^. Branch A (*Islr2 / Cartpt*), Branch B (*Gch1 / Tac1*), and Branch C (*Calcb / Nog*). *May-Zhang et al., 2021 snRNA-seq data only express *Cartpt* at appreciable levels.

### During Early ENS Development the *Sox10^Dom/+^* Allele Alters Cell Allocation Between Two Neuronal Branches

Given previous results showing that the *Sox10^Dom^* allele shifted ratios of enteric neuron classes in the postnatal ENS, we compared the distribution of cells between *Sox10^+/+^* and *Sox10^Dom/+^*at 15.5dpc. We noted that the PC6 cluster is observed exclusively in the *Sox10^Dom/+^* data (Figure 4A). Furthermore, cell quantities are noticeably reduced in neuronal Branch C, while an increase in cell density in neuronal Branch B is apparent in *Sox10^Dom/+^* UMAPs (Figure 4A). We next examined differential cluster abundance to determine whether there were statistically significant differences between *Sox10^+/+^* and *Sox10^Dom/+^*. One-way ANOVA analysis indicated *Sox10^Dom/+^* neuronal populations, cell clusters NC3 (*p* = 0.014) and NC5 (*p* = 0.0378), of Branch C exhibited significant reductions in cell quantities relative to *Sox10^+/+^* replicates. In contrast clusters comprising Branch B (NC2, *p* =.0137; NC0, *p* = .0327) were significantly increased (Figure 4B). We also observed a change in the total cell abundance between genotypes, with the *Sox10^+/+^* data comprised of 3,291 progenitors and 2,292 neuronal cells, while 3,818 progenitors and 1,731 neuronal cells present in the *Sox10^Dom/+^* data.

**Figure 4:**
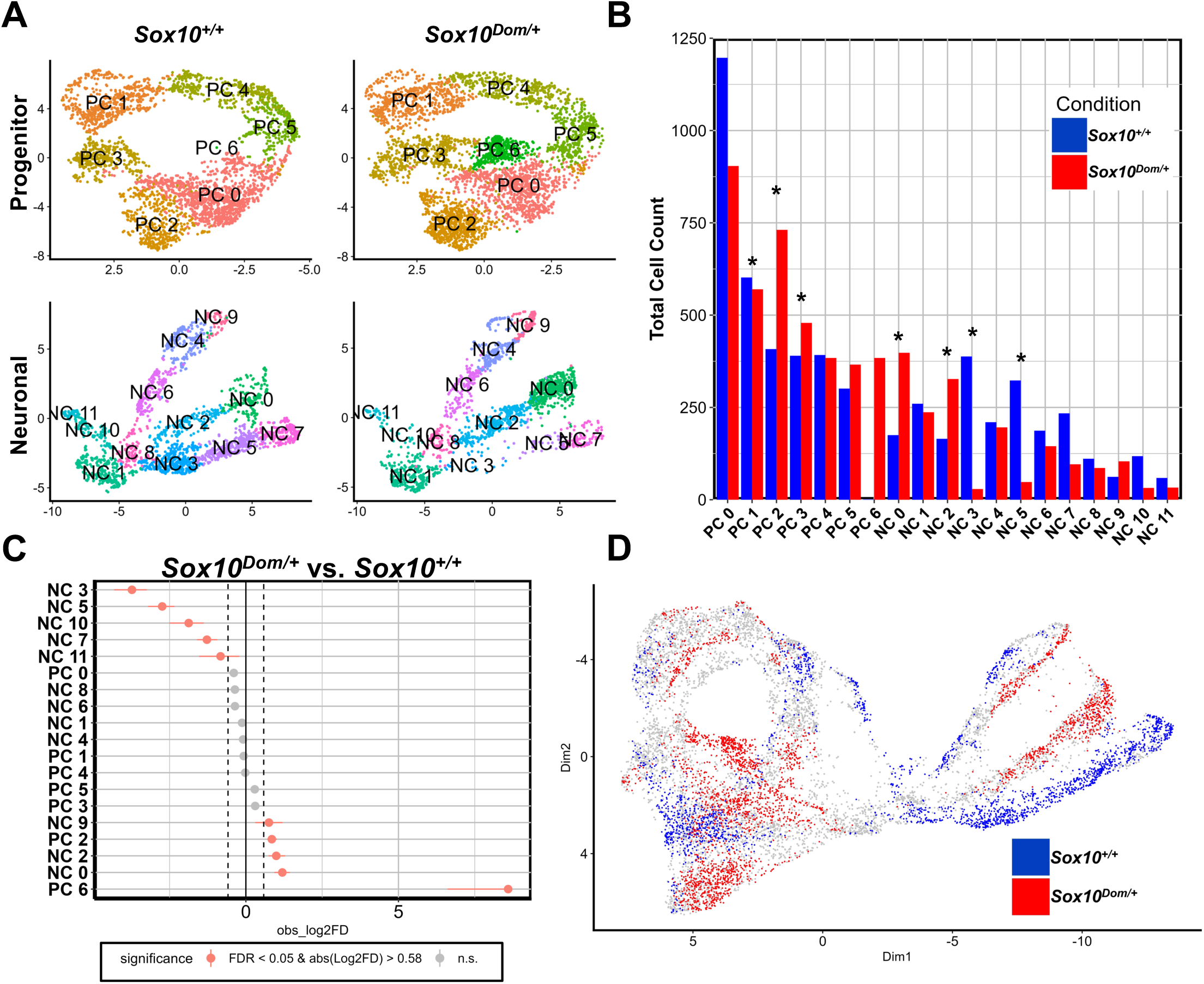
*Sox10^Dom/+^* shifts the distribution of ENS progenitors and differentiating neurons early in ENS development. A) Separate UMAPs of *Sox10^+/+^* and *Sox10^Dom/+^* progenitor and neuronal clusters. B) Bar chart displaying cell counts in every cluster between *Sox10^+/+^* and *Sox10^Dom/+^*lineages. * *p*-value <0.05 from Student’s t-test. C) Relative differences in abundance of cells per cluster between genotypes. Red clusters have an FDR <0.5 and mean Log_2_ fold enrichment >1 (permutation test, n=1,000). Clusters to the left are under-represented in *Sox10^Dom/+^,* while clusters to the right are over-represented in *Sox10^Dom/+^*. D) The DA-seq plot displays a differential abundance of cell populations between genotypes. Cells of increased abundance in *Sox10^+/+^* (blue) compared to those increased in *Sox10^Dom/+^* (red).

To assess variations in cell abundance between genotypes, we utilized additional metrics. Monte Carlo permutation analysis identified a decrease in cell abundance within all clusters associated with Branch C (NC3, NC5, NC7) and an increase in Branch B clusters (NC2, NC0) for the *Sox10^Dom/+^* genotype (Figure 4C). DA-seq analysis was also applied to assess cell abundance irrespective of cluster identities since differences in cell abundance may be attributed to Seurat’s clustering parameters ^29^. In the absence of predefined clusters, DA-seq showed increased cell abundance for *Sox10^Dom/+^* cells along the Branch B trajectory relative to *Sox10^+/+^.* In contrast, *Sox10^+/+^* cells present in Branch C showed a noticeable decrease in cell abundance within the DA-seq UMAP (Figure 4D). Interestingly, the permutation analysis and DA-seq also identified decreased abundance of NC10 and NC11, bridging clusters located between progenitors and neuronal branches, in *Sox10^Dom/+^* cell populations (Figure 4C and 4D). The shifts in cell abundance among transitional bridging populations as well as neuronal branches implicate *Sox10* as a regulator of early neuronal development and enteric neuron diversification.

### *Sox10^Dom/+^* is Associated with Delayed Progression Between Developmental States

Given the abundance changes of transitional bridging clusters (NC10 and NC11) and neuronal branches in *Sox10^Dom/+^* data (Branch B and C), we hypothesized that the *Sox10^Dom^* allele could be altering early cell fate trajectories. To understand how *Sox10* deficits may affect differentiation dynamics of ENCPs, we performed independent RNA velocity analyses for the *Sox10^+/+^* and *Sox10^Dom/+^*datasets using scVelo^30^ (Figure 5A). Two distinct egress regions were observed exiting the main population of cycling progenitors for both genotypes that transition towards neuronal lineages. We designate these two bridge populations Path-1 (NC1, NC8) and Path-2 (NC10, NC11) (Figure 5A). Based on the direction of the velocity plots and position of these cell clusters in the UMAPs, where they reside between progenitors and differentiating neurons, bridge populations Path-1 and Path-2 appear to be transitional in nature. Due to the similarity in the pattern and directionality of cell trajectories between the genotypes, both pathways were examined further to discern any biological characteristics that are distinct between them. Proliferative markers *Miki67, Top2a,* and *Ccnb1* exhibited elevated expression in Path-2 relative to Path-1 for both genotypes (Figure 5B, Table 3). GO analysis was performed on the upregulated genes identified by comparing the two paths (FindAllMarkers, Table 3). Differential gene expression-based Path-2 GO terms were associated with cell replication, whereas Path-1 GO terms were enriched for biological processes linked to neuron maturation and differentiation (DGE Tables 4 and 5). While distinct biological processes were elevated in each pathway, they were found to be comparable between genotypes. Based on these findings, *Sox10^Dom/+^* has minimal effects on overall trajectory patterns as enteric neuronal progenitors transition towards mature fates.

**Figure 5:**
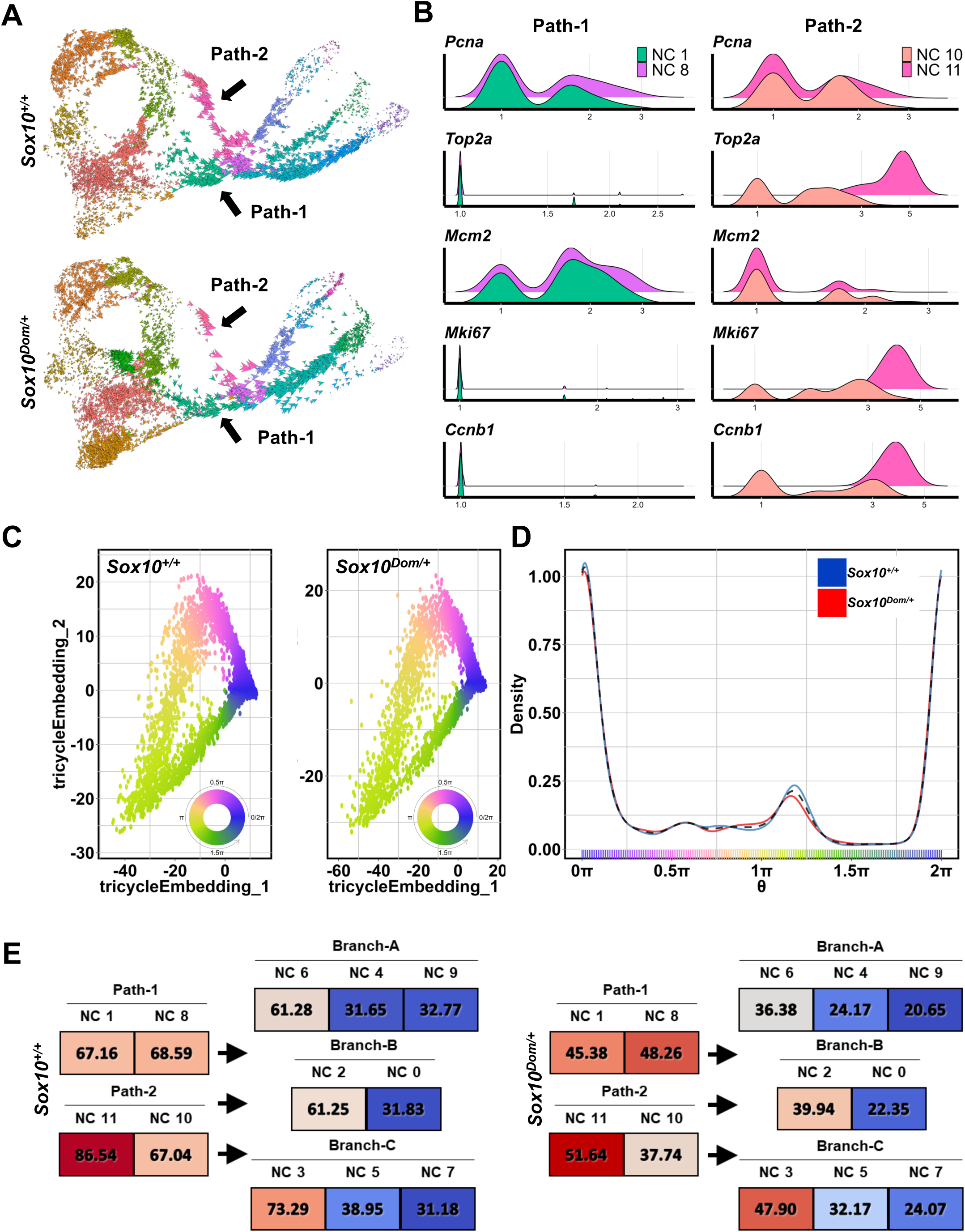
*Sox10^Dom/+^*alters calculated differentiation rates between cell clusters without affecting cell cycle progression. A) RNA Velocity plots show cluster trajectories for *Sox10^+/+^* and *Sox10^Dom/+^.* Two distinct transitional paths, through which progenitors transition towards neuronal fates, are indicated: Path-1 (NC1, NC8) and Path-2 (NC11, NC10). B) Topographical plots depict the expression of proliferative markers *Pcna*, *Top2a*, *Mik67*, and *Ccnb1* along *Ccnb1* transitional Path-1 (NC1, NC8) and Path-2 (NC11, NC10). C) Tricycle analysis embedding distribution of cells in each phase of mitosis for different genotypes. Each annotation corresponds to a specific cell cycle state: 0π (G0/G1), 0.5π (S), 1π (G2), 1.5π (M), and 2π (G0/G1). D) Density plot illustrating the distribution of cells in each phase for *Sox10^+/+^* (blue) and *Sox10^Dom/+^* (red). E) Cell transition rates are calculated via scVelo trajectory analysis for developing ENS lineages of *Sox10^+/+^* and *Sox10^Dom/+^* data. Hues of red indicate higher transition rates, with lower rates represented by blue hues.

Given the role of *Sox10* in melanoma cell cycling^31^ and our observation of a novel progenitor cluster PC6, shifts in cell abundance, and conservation of transition trajectories from progenitors to neuronal branches between genotypes, we examined the association of the *Sox10^Dom/+^* allele with cell cycle progression. To assess whether the phenotypic variations associated with the *Sox10^Dom/+^* allele are linked to enteric neuronal progenitors’ capacity to proliferate and differentiate, each cell was assigned to a cell cycle state via Tricycle analysis^32^. Overall, cell distribution for each cell cycle state remained broadly similar between genotypes, as indicated by the Tricycle embedding patterns and density plots (Figure 5C). Although a slight variation is observed among cells transitioning from the G2 (1π) to the M (1.5π) phase (Figure 5D), the data suggest that cell cycle progression remains consistent and unhindered by the *Sox10^Dom/+^* allele.

As transition trajectories remained consistent and cell cycle progression was comparable between genotypes, we next examined if calculated transition rates are similar along linear trajectories from Path-1 (NC1, NC8) and Path-2 (NC10, NC11) towards the neuronal Branch A (NC6, NC4, NC9), B (NC2, NC0), or C (NC3, NC5, NC7)^30^. The calculated transition rates between genotypes exhibited generally similar patterns, with high rates of transition along Path-1 and Path-2 that gradually decreased as cells progressed towards the more committed neuronal branches (Figure 5E). However, when the calculated transition rates are directly compared between reciprocal clusters in each genotype (e.g., *Sox10^+/+^* NC1 Vs. *Sox10^Dom/+^* NC1), differences in calculated rates of cells moving from one state to the next become apparent (Figure 5E). While cell trajectories and cell cycle progression remain unaffected by the *Sox10^Dom/+^* allele, a general reduction in calculated transition rates along Path-1 and Path-2 suggests that *Sox10* affects neural specification by modulating rates of transition between enteric neuronal progenitor states and neuronal lineages.

### Differential Gene Expression Analysis Identifies Reduced Neurogenic Transcriptional Regulators in *Sox10^Dom/+^* Developing ENS

We next aimed to identify changes in gene expression that could account for the shifts in cellular abundance among enteric neuronal lineages in *Sox10^Dom/+^* mutant data. First, differential gene expression analysis was conducted between genotypes to identify genes that are broadly affected (All *Sox10^+/+^* cells versus all *Sox10^Dom/+^* cells). The analysis identified 1,919 differentially expressed genes (Table 6) of which, 634 were markedly significant (Bonferroni-corrected *p*-value <0.05, average Log_2_-fold change >0.5 or <-0.5). Among the down-regulated genes in *Sox10^Dom/+^*data are multiple factors associated with positive transcriptional regulation of neurogenesis including *Hoxa5*, *Hoxb5*, *Tlx2*, *Hand2os1*, *Hist1h1b*, *Btg1*, *Rasl11a*, and *Phox2b* (Figure 6A). *Pou3f3*, another transcriptional regulator that is important in CNS neuronal development and whose expression is restricted to colonic enteric neurons^3, 33^, was also significantly downregulated in *Sox10^Dom/+^* cells. Differences in gene expression were then compared between genotypes within cell identities (Progenitors, transition pathways, and neuronal branches) to detect lineage and cell type-specific differences. We performed differential gene expression between genotypes per group (Table 7) and filtered for genes with an adjusted *p*-value <0.05 and a Log_2_-fold change of >0.5 or <-0.5. This approach revealed 1,082 unique genes, with several consistently expressed across the developing enteric neuronal lineages. For instance, expression of *Tmsb4x, Uba52, Gm10260, and Gm10076* are increased in all *Sox10^Dom/+^* neuronal branches while *Hoxa5, Hand2os1, and Rbp1* gene expression are decreased (Figure 6C). Despite the overall down-regulation of multiple neurogenic transcription factors (*Phox2b*, *Tlx2*, *Hoxa5*, and *Hoxb5*), we did not observe any lineage specific differences for these well-known regulators of neurogenesis between wildtype and *Sox10^Dom/+^* that could account for the shift in Branch C.

**Figure 6:**
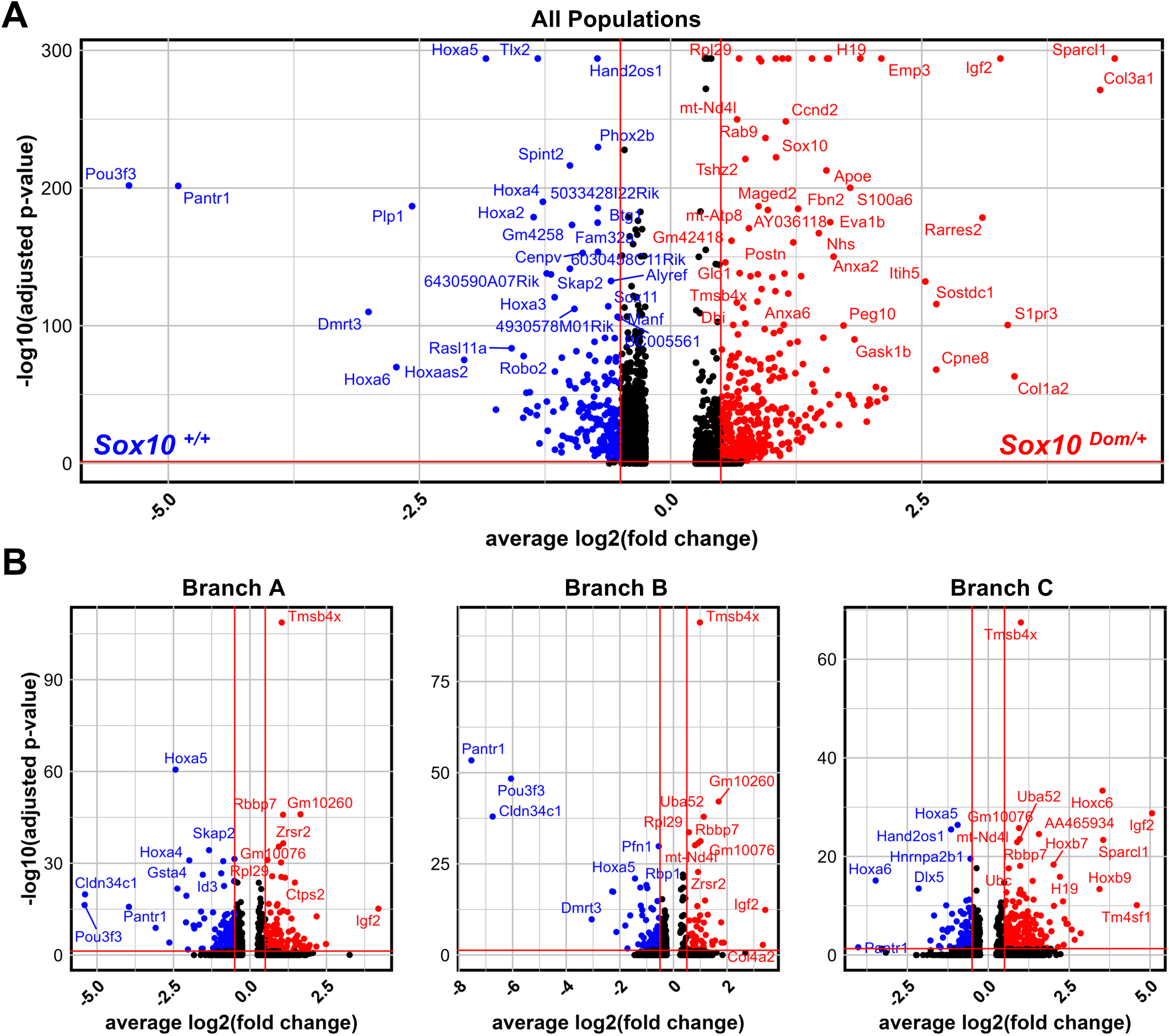
Multiple genes associated with neuronal specification in the developing ENS are significantly downregulated in *Sox10^Dom/+^* lineages. A) Volcano plot displaying the differentially expressed genes identified when comparing all cells from *Sox10^+/+^* and *Sox10^Dom/+^* data. Genes up-regulated in *Sox10^Dom/+^*are in red and genes up-regulated in *Sox10^+/+^* are shown in blue. Genes with an adjusted p-value of 0 were manually changed to be 20-fold less than the lowest non-zero p-value so that the labels for these genes can be shown. B) Individual volcano plots for each neuronal branch within the *Sox10^Dom/+^* data. The genes plotted have an average Log_2_ Fold Change of > 0.5 and < −0.5 and a Bonferroni-adjusted p < 0.05. Down-regulated genes in *Sox10^Dom/+^* are in red while up-regulated genes in *Sox10^Dom/+^* are in blue. Similarly, upregulation in *Sox10^+/+^* is in red while downregulation in *Sox10^+/+^* is in blue.

### Prominent Hox Gene Regulatory Network Activity in ENS Progenitors Transitioning Towards Neuronal Fates

In the absence of discernable differences for well-known regulators of ENS neurogenesis along the *Sox10^Dom/+^* affected neuronal branches, we considered the possibility that the differences in cell abundance within the neuronal trajectories could arise from more subtle shifts in gene regulatory networks (GRN) impacted by the *Sox10^Dom/+^* allele. Initially we sought to identify and derive active gene regulatory networks (GRN) associated with ENS lineages during normal development in *Sox10^+/+^* data. GRNs were assigned using a Single-Cell Regulatory Network Inference and Clustering (SCENIC) analysis^34^. Because SCENIC conducts gene regulatory network reconstruction and cell-state identification by cross-comparing DNA binding motifs and cis-regulatory elements of co-expressed transcripts, it aids in the identification of transcription factors driving cellular heterogeneity. SCENIC analysis identified 141 non-extended regulons, GRNs based on DNA binding motifs and cis-regulatory elements from *Mus musculus* corresponding to the expression of genes in the *Sox10^+/+^* data (Table 8). When GRN activity is observed across all clusters, changes in the relative activity can be tracked as ENS progenitors (PC 0-5) transition along either pathway (NC1, NC8, NC11, and NC10) toward neuronal fates (Figure 7A). Interestingly, we observed a high amount of *Hox* gene associated activity in neuronal branches, with *Hox* GRNs accounting for approximately one-fifth of the active GRNs (9 of 51 total GRNs averaging above a 0.1 activity threshold) in the neuronal Branch A (NC9), B (NC0), and C (NC7) endpoints. In contrast SCENIC analysis in *Sox10^Dom/+^*data showed fewer active *Hox* GRNs overall with only *Hoxa5*, *Hoxb3*, *Hoxb6*, and *Hoxa4* regulons identified (Figure 7B).

**Figure 7:**
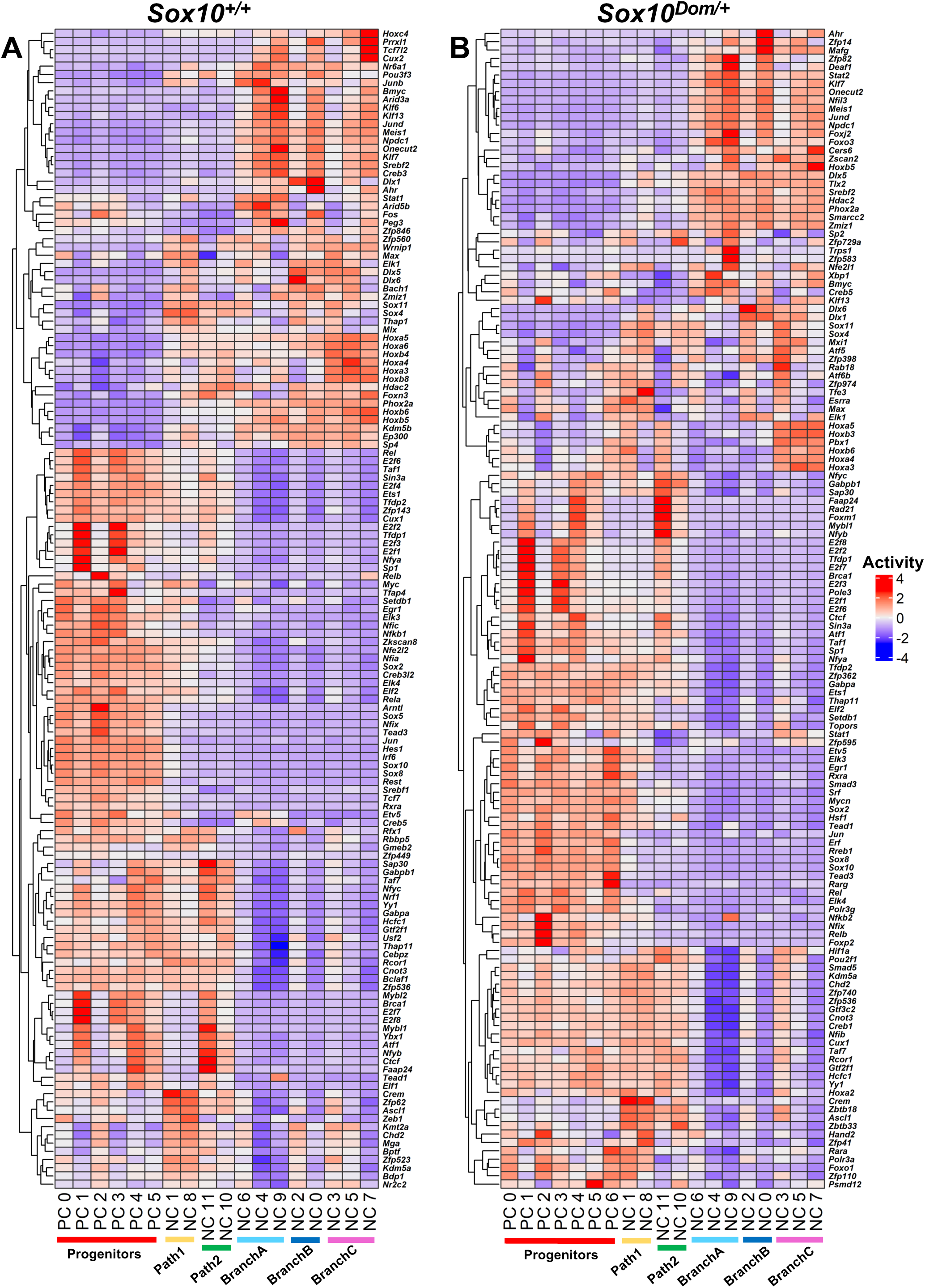
Multiple gene regulatory networks that exhibit elevated activity within developing enteric neuronal lineages are associated with Hox-related genes. Heatmap of gene regulatory network activity calculated via SCENIC for wildtype *Sox10^+/+^* (A) and *Sox10^Dom/+^* (B) data. Elevated activity is shown as hues of red with decreased activity indicated by hues of blue for each gene-associated regulon.

### *Hoxa6* is Downregulated in Developing *Sox10^Dom/+^* Enteric Neuronal Lineages

Having identified the prominence of Hox GRN via SCENIC in wild type samples and the reduced presence of Hox-associated regulons in *Sox10^Dom/+^*, we performed a focused assessment of *Hox* genes in *Sox10^Dom/+^*data compared to *Sox10^+/+^*. We observed that *Hoxa3*, *Hoxa4*, *Hoxa6*, and *Hoxb6* are dysregulated in distinct *Sox10^Dom/+^*clusters (Figure 8A). Interestingly in wild-type data, we found that *Hoxa6* expression is normally elevated in Path-2 and Branch C, the populations that are significantly affected in *Sox10^Dom/+^* mutants. Moreover, in *Sox10^Dom/+^*mutant data, *Hoxa6* is significantly downregulated in Path-2 and Branch C, most notably in cells among neuronal clusters (Figure 8B; Table 7).

**Figure 8:**
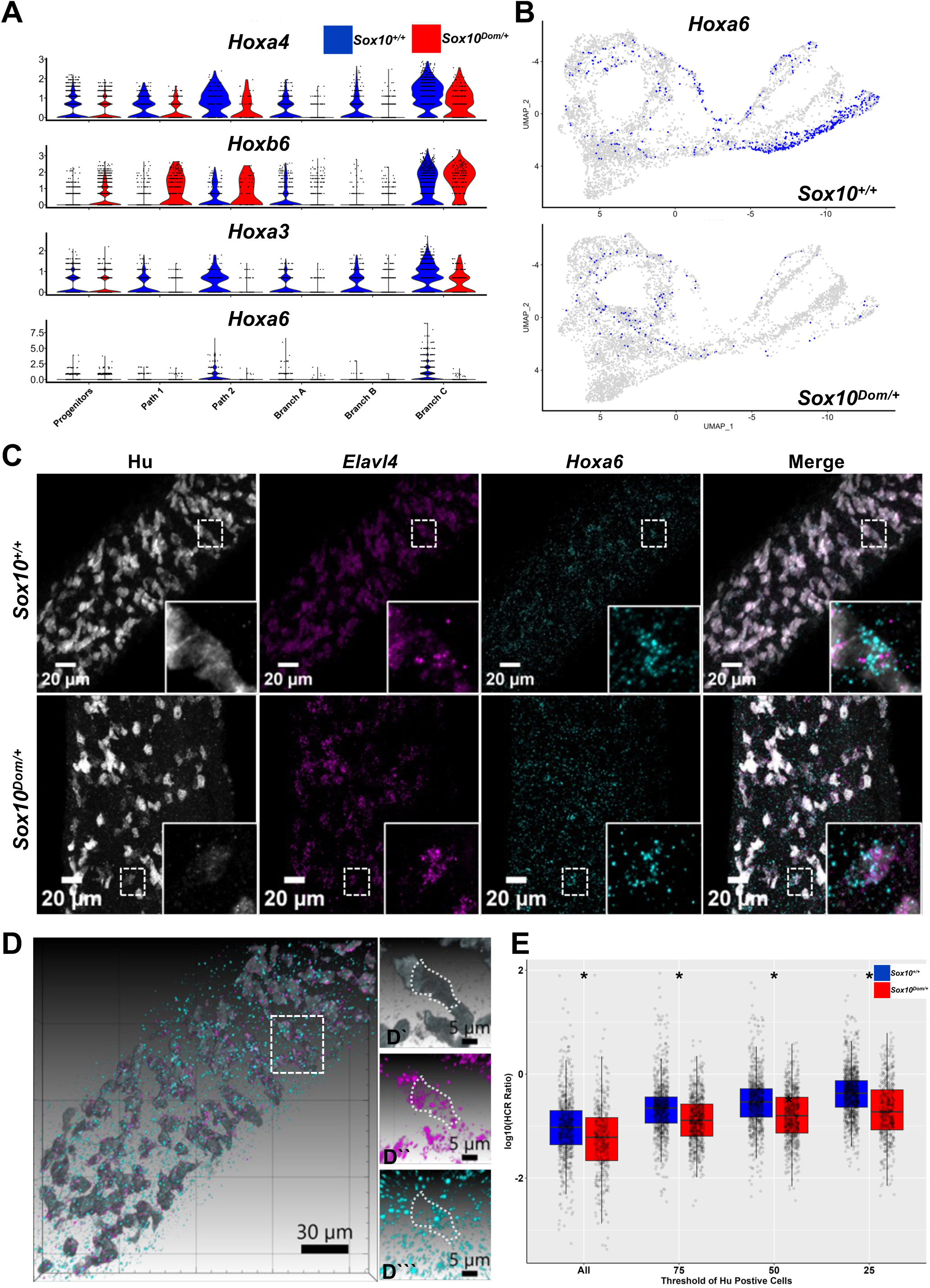
*Hox* transcripts associated with active regulatory networks are differentially expressed between *Sox10^+/+^* and *Sox10^Dom/+^* ENS populations. A) Violin plots show the normalized Log_2_-fold expression levels of *Hox* genes in *Sox10^+/+^*(blue) versus *Sox10^Dom/+^* (red) cells. B) Feature plot UMAPs depict *Hoxa6* expression patterns in *Sox10^+/+^* and *Sox10^Dom/+^*. C) Fluorescent confocal images of HCR *in situ* stained for HuC/D in 15.5 dpc ileum from *Sox10^+/+^* and *Sox10^Dom/+^* fetal mice. Insets emphasize intensity of *Hoxa6* HCR signal in dim HuC/D+ cells D) Images from 3D rendering highlight the spatial localization of *Hoxa6* in dim Hu C/D+ cells within a *Sox10^+/+^* sample (Refer to videos 1 and 2) E) Boxplot displaying positive, Log_10_-transformed HCR signal ratios (*Hoxa6* / *Elavl4*) in wholemount fetal ileum samples based on all HuC/D+ cells (100%) to the dimmest quartile of HuC/D+ cell area (25%). Asterisk indicates Wilcoxon non-paired test *p*-values between genotype (*p*< 0.05). Each datapoint represents HCR signal ratio of one region of interest in a given image.

We applied hybridization chain reaction (HCR) coupled with immunolabeling of developing neurons using the pan-neuronal marker HuC/D to validate changes in *Hoxa6* expression *in situ*. HCR Probes were used to localize mRNA transcripts for the gene *Elavl4* (transcript for HuD) in parallel with *Hoxa6.* HuC/D immunolabeling was applied to *Sox10^+/+^* and *Sox10^Dom/+^* fetal intestines at 15.5 dpc to localize emerging enteric neurons that express this antigen. We concentrated our imaging analysis on the proximal ileum to detect differentiating neurons in *Sox10^Dom/+^* mutants that exhibit variable aganglionosis in the colon^22^. To ensure samples exhibited the same degree of phenotype severity as those processed for scRNA-seq, we selectively processed samples with absence of ENS cells distal to the cecum based on *Phox2b*-CFP transgene expression. After HCR and immunolabeling, colocalization of the *Hoxa6* signal with *Elavl4* HCR probe signal in HuC/D+ cells in the ileum for both *Sox10^+/+^* and *Sox10^Dom/+^* fetal intestines was observed (Figure 8C). Interestingly, while imaging biological replicates for both *Sox10^+/+^* and *Sox10^Dom/+^*fetal intestines, we noted greater *Hoxa6* signal intensity in dimly labeled HuC/D+ cells that was particularly prominent in *Sox10^Dom/+^* samples. Three-dimensional renderings improve visualization of the higher intensity of *Hoxa6* transcript in these dim HuC/D+ cells (Figure 8D, Videos 1-2). Consistent with the single-cell observations, ratios of HCR signal intensity for *Hoxa6* relative to *Elavl4* signal between *Sox10^+/+^* and *Sox10^Dom/+^* genotypes revealed a significant difference in the *Hoxa6* mRNA levels.

To focus our analysis on the population of dim HuC/D cells and facilitate detection of differences between genotypes, we systematically applied thresholds for HuC/D+ cells based on signal intensity. Starting from the baseline of 100% H C/D signal intensity, thresholds were set to capture signal for *Hoxa6* within the 75% of the least dim HuC/D+ cell population, 50% of the least dim HuC/D+ cells, and the 25% most dimly labeled HuC/D+ cell populations. Selective quantification of dimly labeled HuC/D+ cells revealed elevated ratios of *Hoxa6* HCR signal relative to *Elavl4* between genotypes as thresholding became more restrictive (Figure 8E). Comparison of *Hoxa6* / *Elavl4* HCR ratios based on intensity thresholds between *Sox10^+/+^* versus *Sox10^Dom/+^* genotype revealed a significant difference for all HuC/D+ cells (*p* = 1.46e-08), the 75% dimmest (*p* = 2.20e-19), and most prominently in the 50% (*p* = 4.82e-23) and 25% dimmest HuC/D+ cells (*p* = 1.06e-29). Since dysregulation of *Hoxa6* correlates with a shift in Branch C abundance in *Sox10^Dom/+^* single-cell data and *Hoxa6* is differentially expressed between genotypes, most notably in dim HuC/D+ cells that are initiating neurogenic processes, this gene emerges as a logical candidate regulator of enteric neuron specification in ENS development. A scan of the *Hoxa6* locus for SOX10 binding motifs revealed multiple consensus binding motifs within the *Hoxa6* locus that further strengthens the likelihood of interaction between these two genes.

## Discussion

Recent single-cell studies have dramatically expanded our understanding of cellular diversity within the developing and mature ENS^2-4, 33, 35^. Each effort to map ENS development has identified consistent populations of progenitors and diverse neuronal trajectories bridged by transitioning neuroblasts^4, 17, 36-38^. However, understanding of the factors that regulate neuronal diversity in the ENS has been limited due to the spatial complexity of neuronal maturation along the fetal intestine with neurons maturing behind the initial wave of progenitors colonizing the fetal bowel. As a result, relatively few factors that regulate diversification of specific enteric neuron types have been identified^4, 18^. Our study significantly broadens understanding of factors regulating early enteric neuron diversity through analysis of the *Sox10^Dom/+^*allele. This work has revealed three distinct neuronal trajectories during early ENS lineage divergence. In *Sox10^Dom/+^* mutants, two of these neuronal trajectories are shifted in cellular abundance. We found that multiple transcriptional regulators that are positively associated with neurogenesis are reduced in *Sox10^Dom/+^* mutants. Moreover, our analysis identified multiple *Hox* genes with increased GRN activity during enteric neurogenesis in wild-type cells of which several were dysregulated in *Sox10^Dom/+^* ENS populations. Most prominently, we found down-regulation of *Hoxa6* in early differentiating neurons that emphasizes this factor as a logical candidate in specification of enteric neuron subtypes. Our findings are consistent with the hypothesis that dysregulation of *Sox10* disrupts GRNs in ENS progenitors and feeds forward to alter essential effector genes in enteric neuron diversification. Collectively, the results illustrate the value of leveraging mutant alleles, like *Sox10^Dom/+^*, to elucidate regulatory factors in the gene networks that control ENS neurogenesis.

The three distinct neuronal branches captured in our analysis at 15.5dpc likely results from utilization of the *Phox2b*-CFP transgene. This line comprehensively labels all enteric neurons in both small intestine and colon and overlaps completely with PHOX2B protein immunolabeling^20^. Other ENS scRNA-seq studies have relied on combinatorial cre-loxP approaches that require activation of a fluorescent reporter for cell labeling^4, 33^. In our hands, the traditional neural crest fate mapping line, *Wnt1*-cre (H2az2^Tg(Wnt1-cre)11Rth^, RRID: MGI:238657) does not label all enteric neurons in the postnatal ENS^19^. Others have similarly published incomplete coincidence of cre reporters with labeling of postnatal enteric neurons using *Wnt1*-cre*, BAF53b-*cre, or *Sox10*-creER^39, 40^. Another unlikely possibility is that the third neuronal branch could have arisen from differences in analysis methods between studies. However, even after comparable batch correction and integration of our data with that of Morarach and colleagues for direct comparison, the third distinct neuronal branch remained (Figure 2C). Thus, the third neuronal trajectory most likely arises from inherent differences in the dataset either from the specific mouse lines used or the techniques of isolation. Despite the difference in number of distinct neuronal branches profiled, our studies are consistent with those of Morarach and colleagues who reported postmitotic neuronal diversification after ENCPs take on neuroblast identity^4^.

Capturing the full diversity of enteric neurons and developing markers to distinguish neuron subtypes has been an important goal in enteric neurobiology. However, the ability to track development of discrete neuron types over developmental time is still an open area of investigation that would benefit from identification of new marker genes for emerging neuronal subtypes. To identify candidate markers that might be useful for future lineage tracing we examined expression of genes in cells at the furthest tips of the three neuronal lineages in our 15.5dpc data across other more mature datasets. We found that these marker sets (Branch A, *Islr2 + Cartpt*; Branch B, *Tac1+Mdga1*; Branch C, *Nog+Calcb*) are consistently present in distinct populations of cells across multiple different datasets. This observation suggests that the expression of these genes or reporters driven from their regulatory regions may be used in future studies to track neurons within the distinct trajectories as they proceed through subsequent maturation.

ENS development is complex and there are two obvious possible explanations for the effect of *Sox10^Dom/+^*on mature enteric neuron ratios. Aberrant enteric glial development could be altering enteric neuron differentiation. Alternatively, the *Sox10^Dom^*mutation in ENCPs may be feeding forward to alter essential GRNs needed for enteric neuron diversification. Our findings are consistent with an effect of the *Sox10^Dom/+^* allele in progenitors altering enteric neurogenesis despite absence of SOX10 in differentiating enteric neurons. Differential expression of several genes associated with transcriptional regulation of neurogenesis in *Sox10^Dom/+^* mutants likely explains how deficits in *Sox10* lead to alterations in enteric neuron ratios. Multiple transcription factors that have been implicated in ENS development are differentially expressed in *Sox10^Dom/+^* populations compared to wildtype including *Phox2b*, *Hoxb5*, *Tlx2*, and *Zfhx3.* Deficits of enteric ganglia are well known for mutations in *Phox2b* (Congenital Central Hypoventilation Syndrome with aganglionosis)^41^ and *Tlx2* (neuronal intestinal dysplasia)^42^, while *Zfhx3* is associated with HSCR^43^. In zebrafish, *Hoxb5b* has been implicated in expansion of ENS progenitors and overexpression of this gene reduces neurogenesis, emphasizing the critical interplay between expression level and developmental progression in the ENS^44^. We also observed absence of *Pou3f3* in *Sox10^Dom/+^* mutants. *Pou3f3* is present in colonic neurons^3, 33, 45^ and the lack of expression in our *Sox10^Dom/+^*data is likely due to our selection of severe *Sox10^Dom/+^* mutants that lacked ENS in the colon for our profiling studies. While certainly related to broad neurogenesis, disruption of these genes does not explain the effect of the *Sox10^Dom/+^* allele on specific branches of developing enteric neurons.

It is intriguing that the impact of *Sox10^Dom/+^* genotype on neuronal allocation is not uniform across all lineages, despite *Sox10* being widely expressed in ENS progenitors. Relatively few genes have been identified that exert comparable effects in disrupting the abundance of specific enteric neuron subtypes. Loss of *Ascl1* leads to decreased *Vip* and *Calbindin*-expressing neurons (Branch A derived neuron subtypes)^46^. Similarly, deletion of *Pbx3* leads to fewer CALB+ neurons in the Branch A trajectory^4^. In our *Sox10^Dom/+^* datasets we see no decrease in *Vip* and *Calbindin*-expressing cells that arise in Branch A. One distinction between our results and those found for *Ascl1* and *Pbx3* is that both *Ascl1* and *Pbx3* are expressed in developing neurons, while *Sox10* is present in progenitors and is extinguished in developing neurons. We did find that the long non-coding RNA *Hand2os1*, a known regulator of *Hand2 ^47^,* as well as *Hand2* itself, which plays a central role in specification of enteric neurons^48, 49^, are both downregulated in *Sox10^Dom/+^* progenitor and neuronal clusters (Table 7). However down-regulation of *Hand2* also does not account for the branch specific shifts in neuron types observed in *Sox10^Dom/+^* mutants as haploinsufficiency of *Hand2* leads to broader deficiency of enteric neuron subtypes including both NOS1+ and Calretinin+ neurons^48, 49^. Thus, the factors mediating the shift in neuronal allocation between the enteric neuronal branches B and C remain to be identified and will be essential for stem cell engineering efforts that seek to replace specific neuronal lineages in the treatment of GI motility disorders.

Our analysis specifically emphasizes *Hox* gene activity in ENS populations undergoing neurogenesis. We found that several *Hox* genes are differentially expressed in our comparisons of *Sox10^+/+^* and *Sox10^Dom/+^*data. GRN analysis from SCENIC in the *Sox10^+/+^* data revealed that nearly a fifth of regulons within neuronal populations are linked to Hox genes, with increased activity of *Hoxa3, a4, a5, a6, b4, b5, b6, b8, and c4* in transitioning neuroblast and neuronal clusters (Figure 7). *Hox* genes are well known for their involvement in patterning the brain and spinal cord during development and exhibit analogous regional expression in the gut that impacts gross anatomy of the developing intestine^50^. Prior work had suggested *Hox* genes could be involved in ENS neurogenesis and *Hox* mutations or altered expression are associated with megacolon phenotypes in mice and patients ^51-56^. In zebrafish a prominent role for *Hoxb5b* has been identified as a regulator governing the proliferation of vagal ENS progenitors and appropriate progression of these progenitors broadly into neurons^44^. Of the *Hox* genes identified by GRN analysis, *Hoxa6* expression is prominently localized within cells along Path-2 and Branch C of *Sox10^+/+^*data, the populations of cells most affected in the *Sox10^Dom/+^* mutants. Moreover, *Hoxa6* is significantly downregulated in the *Sox10^Dom/+^* data for these groups. The heightened activity of the *Hoxa6* regulon during normally developing neuronal lineages combined with the prominent *Hoxa6* down-regulation in Branch C *Sox10^Dom/+^* data suggest this factor as a relevant regulator of enteric neuron subtype specification.

Both scRNA-seq and *in situ* analyses indicate that *Hoxa6* expression is altered in ENS populations of *Sox10^Dom/+^* mutants. By implementing HCR for *in situ* localization, which enables single cell visualization, we observed higher *Hoxa6* HCR ratios (*Hoxa6*/*Elavl4* HCR signal) in emerging neurons that were weakly positive for HuC/D protein in wild-type samples. In contrast, in the *Sox10^Dom/+^* fetal gut we observed a significant decrease in *Hoxa6/Elavl4* HCR signal ratios compared to *Sox10^+/+^*ratios. A well-known marker of developing neurons, low levels of HuC/D pinpoints early emerging neurons that enables stratification of developing neurons based on HuC/D signal intensity. Our observations suggest a correlation between the decline in Path 2 and Branch C enteric neurons and the reduction of *Hoxa6* positive cells. Whether the regulation of *Hoxa6* in ENS progenitors is directly mediated by SOX10 remains to be seen. Previous chromatin immunoprecipitation studies in rat spinal cord suggest that SOX10 can directly interact with the promoter region of *Hoxa6^57^* and we identify SOX10 consensus binding motifs within the mouse *Hoxa6* locus. Future studies that build on our identification of *Hoxa6* and other *Hox* genes will be needed to fully define the GRNs regulating allocation of enteric neuron subtypes.

Whether *Sox10* mediates its effects on enteric neuron lineages via direct regulation of downstream transcription factors or is more broadly regulating chromatin accessibility is an area for future study. We have identified SOX10 consensus binding motifs in the Hoxa6 locus that suggest this gene is a direct target. However, *Sox10* effects on ENS development are likely multimodal. *Sox10* has been implicated in regulation of chromatin accessibility in both melanocyte and Schwann cell differentiation^58-60^. *Sox10* physically interacts with the BAF chromatin-remodeling complex to activate Schwann cell differentiation programs^58^. Given the multifunctional nature of *Sox10* effects in other aspects of neural crest development, the gene may be exerting both gene-specific direct regulation via promoter activation as well as exerting a more diffuse effect on overall chromatin accessibility. Future work to localize *Sox10* direct targets in ENS progenitors is needed to elucidate primary nodes in GRNs versus those that result from SOX10 effects on chromatin accessibility.

The data presented in this study refines and expands the framework of neuronal diversification during ENS development (Figure 9). Three neuronal trajectories arise as ENCPs differentiate along two transitional pathways. These transitional paths are distinguished by differences in proliferative capacity. The data shows that *Sox10* affects enteric neuron subtype allocation early in ENS formation along two of the three neuronal trajectories. The change in enteric neuron subtype distribution coincides with the downregulation of positive transcriptional regulators and a reduction in the calculated rate of progenitor transition to neuronal states. Finally, we detect elevated *Hox* GRN activity during enteric neurogenesis, with several *Hox* genes exhibiting dysregulation in the context of the *Sox10^Dom/+^* allele, including the Branch C lineage specific *Hoxa6*. Our analysis implicates *Sox10* as multifunctional factor that, in addition to its other roles in neural crest development, also participates in early enteric neuronal diversification in the developing ENS.

**Figure 9:**
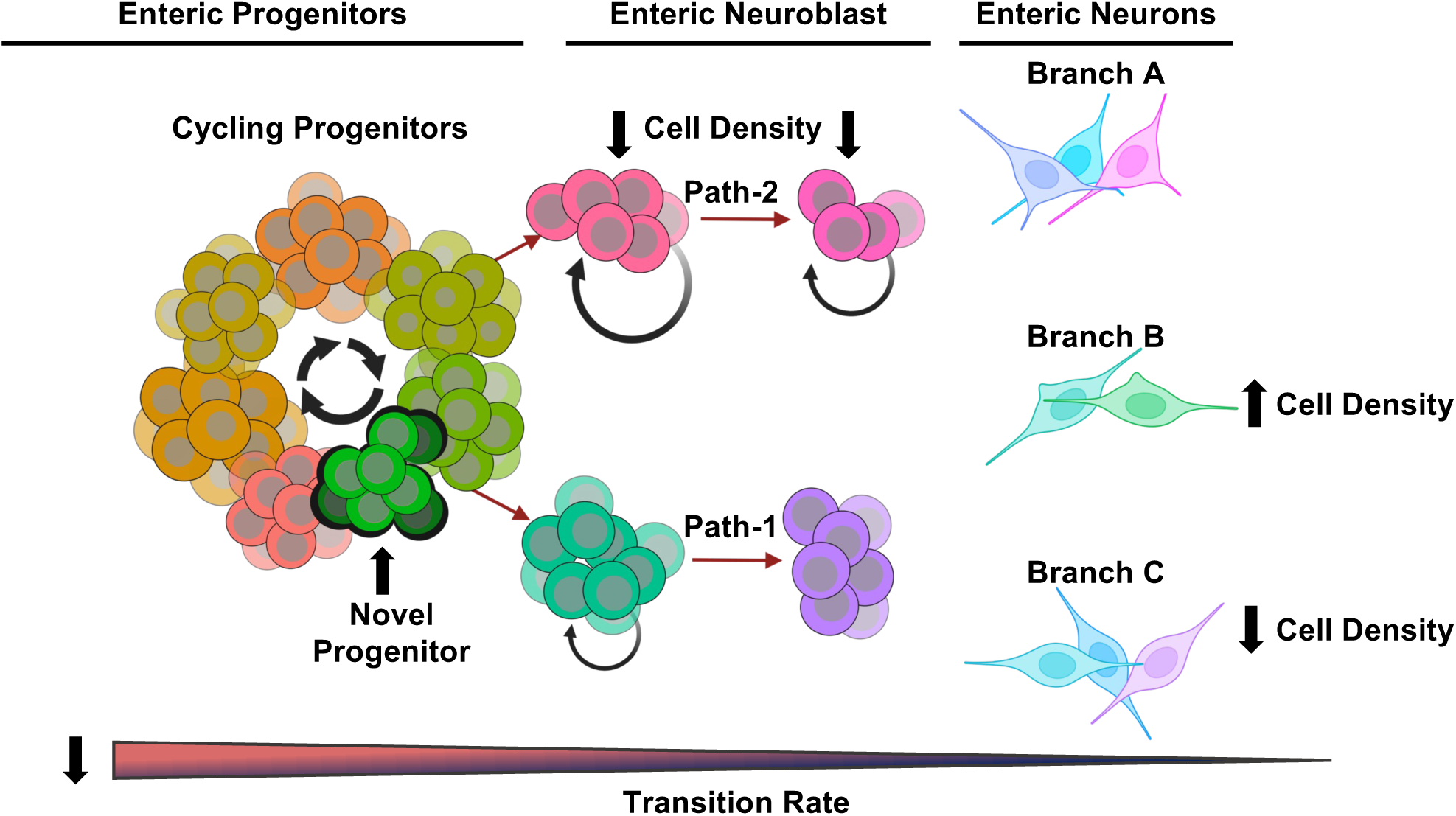
Observations and Model of *Sox10*-Mediated Enteric Neurogenesis. As early as 15.5 dpc, distinct ENS progenitors, two neuroblast transition pathways, and three distinct neuronal lineages can be identified via scRNA-seq. Perturbation of ENS development by the *Sox10^Dom/+^* mutant allele produces a novel progenitor population and causes a noticeable shift in lineage distribution associated with decreased cell abundance within Branch C and Path 2, and an increase in Branch B (bold arrows). Downregulation of genes associated with neuronal specification are detected, likely contributing to the calculated decrease in transition rates between cell states.

### Summary

Treatment of enteric neuropathies remains a major challenge for gastroenterologists. Lack of knowledge regarding the regulatory factors that control diversification of ENS cell types has inhibited directed differentiation of specific enteric neuron subtypes. Here, we applied single-cell RNA-Seq to demonstrate new roles for *Sox10* in early enteric neuron lineage allocation and identified multiple GRNs, including several *Hox* regulons, in differentiating enteric neurons. Prominent expression of *Hoxa6* in populations affected by *Sox10* mutation and coincident down-regulation of *Hoxa6* in *Sox10^Dom/+^* mutants implicates *Hoxa6* in specification of some enteric neuron types.

## Methods

### Animals

The Institutional Animal Care and Use Committee at Vanderbilt University approved all experimental protocols. All mice were housed in a modified barrier facility on a 14-hour on, 10-hour off-light cycle in high-density caging (Lab Products Inc., #10025) with a standard diet (Purina Diet #5L0D) and water ad libitum. Mice carrying the Dominant megacolon (*Sox10^Dom^*; RRID: MGI:1857401), hereafter referred to as *Sox10^Dom/+^*, were maintained as congenic stocks by backcrosses with C57BL/6J (Jackson Laboratory Stock# 664) for more than 20 generations. Timed mattings were set to obtain staged mouse fetuses, designating the morning of plug formation as 0.5 days post coitus (dpc). The staging of fetuses was confirmed by examination of the fore and hind limbs compared to published standards^61^. Genotyping was performed using primers for the *Sox10^Dom^* allele as previously described^9^. The Tg(Phox2b-HIST2H2BE/Cerulean)1Sout mice (MGI: 5013571) and Tg(Sox10-HIST2H2BE/YFP*)1Sout mice (MGI:3769269), referred to as *Phox2b*-CFP and *Sox10*-YFP mice, were crossed with *Sox10^Dom/+^* mice to generate *Phox2b*-CFP / *Sox10*-YFP / *Sox10^Dom/+^* and *Phox2b*-CFP / *Sox10*-YFP / *Sox10^+/+^* offspring.

### Isolation of ENS Progenitors

Mouse fetuses at 15.5 dpc were screened for expression of CFP and YFP transgenes using a Leica 205FA fluorescent stereomicroscope. Fetal gastrointestinal tracts, extending from the stomach to the anus, were dissected from embryos that exhibited dual-positive fluorescent transgene expression. The extent of aganglionosis based on transgene fluorescence in the hindgut was assessed for each fetal intestine, and individual samples were processed separately. Fetal intestines were dissociated in a 1 ml solution containing Native Bacillus Licheniformis (Creative Enzymes: NATE-0633) at a concentration of 5 mg/ml, diluted in HBSS (Gibco:14185-052), supplemented with 30 μl of DNase I (Sigma:D4527) and incubated in a 5°C water bath. Subsequently, the dissociated cells were filtered multiple times through a 40-μm nylon strainer into 5 mL round-bottom collection tubes. Dissociated enteric lineages were rinsed with L-15 medium (Gibco: 21083-027), consisting of penicillin/streptomycin (Gibco:15140122), amphotericin (Corning MT30003CF), BSA (Sigma:A3912), HEPES and BioWhittaker water(Fisher: BW17-724Q). A Viability dye (7-AAD, Invitrogen: A1310) at a 1:1000 dilution was added to cell suspension before cell isolation to exclude nonviable cells. Following filtration, all cells exhibiting CFP or YFP fluorescence were isolated via flow sorting using a 5-Laser FACS Aria lll equipped with a 100-micron nozzle into RNase-free PBS with 0.5% BSA. *Phox2b-CFP* and *Sox10-YFP* expressing cells were identified through a series of gating procedures, ensuring the isolation of single viable cells positive for the expression of these fluorescent transgenes.

### Generation of single cell libraries

Single cells flow sorted at 15.5 dpc were encapsulated using 10X Genomics 3.1 Chromium Single Cell 3’ GEM library reagents. Libraries were sequenced using a Nova-seq 6000 to produce paired-end 50-bp sequencing reads. Four *Sox10^+/+^* biological replicates and three *Sox10^Dom/+^* replicates were generated. Below is a summary of sample data generated by 10x Cell Ranger 5.0 using reference sequence mm10-2020-A.

| **Sample** | **Estimated Number of Cells** | **Mean Reads Per Cell** | **Mean Genes Per Cell** | **Total Reads** |
| --- | --- | --- | --- | --- |
| Sox10<sup>+/+</sup> R1 | 3,538 | 96,052 | 1,784 | 339,832,005 |
| Sox10<sup>+/+</sup> R2 | 2,947 | 129,880 | 4,661 | 382,756,532 |
| Sox10<sup>+/+</sup> R3 | 3,469 | 94,771 | 2,521 | 328,760,675 |
| Sox10<sup>+/+</sup> R4 | 1,427 | 374,895 | 4,896 | 534,974,456 |
| Sox10<sup>Dom/+</sup> R1 | 4,798 | 78,353 | 3,388 | 375,938,797 |
| Sox10<sup>Dom/+</sup> R2 | 4,818 | 105,119 | 3,929 | 506,461,671 |
| Sox10<sup>Dom/+</sup> R3 | 5,087 | 72,885 | 4,288 | 370,767,155 |

### Data Analysis

Default settings for all software and packages were used unless stated otherwise. Data was initially analyzed via the 10x Cell Ranger 3.1 using reference sequence mm10-2020-A. The raw and filtered CellRanger outputs were imported via SoupX v1.5.2 to profile and remove ambient mRNA contamination in R v4.1.2 ^62^. Runs from the *Sox10^+/+^* and *Sox10^Dom/+^* were separately integrated and batch-corrected using the Seurat SCT integration (v4.1.1), and cells were selected based on a minimum of 200 features per cell and the exclusion of cells with more than 10% mitochondrial gene expression^63^. After, *Sox10^+/+^*and *Sox10^Dom/+^* data were merged, SCT data and preexisting variable features were maintained. Subsequent, unsupervised Louvain clustering and UMAP analysis were performed using the SCT assay within the merged data with “FindNeighbors” set to 1:30 dimensions and “FindClusters” resolution set to 0.5. Initial cell identities and group distributions were assigned based on known ENS progenitor and neuronal markers. Gene lists were generated form *Sox10^+/+^* data using Seurat’s “FindAllMarkers” analysis with the Wilcoxon rank-sum test and Bonferroni p-value correction (p <0.05) using Seurat v5.0.

Gene ontology analysis was performed using ClusterProfiler with input of gene lists identified from “FindAllMarkers” analysis (Table 2) to determine the biological processes linked to each group and establish cluster identities^64^. Differentially expressed genes (Bonferroni-adjusted p-value <0.05, average Log_2_(FoldChange) > 0) from FindAllMarkers results for each of the three main groups’ (Progenitors, Neuronal Branches, Mixed/Unknown) were used as input for MGI Batch search (https://www.informatics.jax.org/batch_data.shtml, August 2024)^65^ to obtain Entrez IDs for input for clusterProfiler’s enrichGO function. The same methods were applied to the differentially expressed genes for the Path1 and Path2 clusters. High ribosomal gene expression, high mitochondrial and low RNA feature counts led to exclusion of clusters 1, 10, and 14 from further analysis^66, 67^. Subsequently progenitor and neuronal linages were separated, and the SCT assay was used with “FindNeighbors” set to 1:20 dimensions and “FindClusters” resolution set at 0.5 to identify sub clusters. The newly formed clusters were labeled as Progenitor Clusters (PC) and Neuronal Clusters (NC) and stored in the metadata of the combined progenitor and neuronal object. The embryonic datasets from Morarach and colleagues were processed using comparable techniques^4^. Differential expression analysis was performed using Seurat’s “FindAllMarkers” function in Seurat V5.0. The analysis used a minimum percentage (min.pct) of 0.1 and an average Log_2_-fold change threshold of >0.25 and <-0.25 to identify differently expressed genes. Genes with Bonferroni corrected *p*-value <0.05 and a Log_2_-fold change of of >0.5 and <-0.5 were considered differentially expressed.

Harmony Batch correction was utilized to identify conserved cell types within the age matched *Sox10^+/+^* and Morarach 15.5 dpc datasets^25^. The default normalization parameters for Harmony reduction and integration were utilized to generate Harmony embeddings. Post batch correction, the Harmony assay was used with “FindNeighbors” set to 1:20 dimensions and “FindClusters” resolution set at 0.5 to generate UMAPs of the batch-corrected datasets.

A computational method for single-cell regulatory network inference and clustering (SCENIC) analysis was used to identify gene regulatory networks and their corresponding cellular activity using pySCENIC version 1.2.4^34^. SCENIC provides a database of cis-regulatory element transcription factor binding motifs that are over-represented in over 30 cis-regulatory element datasets identified using RcisTarget^34^. A database of 20,000 motifs obtained from RcisTarget and GENIE3^68^ was utilized to establish transcript regulons. Gene regulatory networks were identified using the expression matrix from *Sox10^+/+^*and *Sox10^Dom/+^* separately.

To predict cell trajectory and transition rates, ScVelo V0.3.2 was employed. Spliced and unspliced reads were annotated using Velocyto^30, 69^. Splice ratios for *Sox10^+/+^* were 77% spliced and 23% unspliced while *Sox10^Dom/+^* ratios were 76% spliced and 24% unspliced. UMAPs from Seurat were split for ease of interpretation. The calculated velocity vectors were then compared between the genotypes to evaluate differences in the differentiation rates.

Cell cycle annotation and quantification were performed using Tricycle V1.8.0^32^. Expression of genes associated with cell cycling, such as *Top2a* and *Smc2*, were examined to verify the cell cycle annotation. The density plot generated from the Tricycle analysis employed a von Mises distribution, mapping the phases of the cell cycle in a circular fashion: 0π (G0/G1), 0.5π (S), 1π (G2), 1.5π (M), and 2π (G0/G1).

The difference in proportions of cells within clusters between genotypes was analyzed using scProportionTest V 0.0.0.9 ^70^. A permutation test was employed to calculate a *p*-value for each cluster, assessing the significance of the observed differences in proportions between genotypes. Additionally, a confidence interval for the magnitude of the difference was determined via 1000 bootstrapped iterations.

Differential abundance (DA) analysis was conducted to examine variations in cell distribution by grouping cells based on their k-nearest neighbors. This approach allows for observing differences that are not constrained by predefined clusters. DA-seq version 1.0.0 was utilized for this analysis^29^. The DA-seq plot illustrates the differential abundance of cell populations between genotypes, determined based on vector scores. These scores were further subjected to random permutation to generate the null distribution for the DA measure accurately. The upper and lower thresholds, set to ±0.8 of increased and decreased DA measurement values, respectively, ensure cells exceeding these limits are not false positives and predict differentially abundant cells within the data.

*Hoxa6* locus Sox10 binding motifs were detected via scanning the mm10 genome for all possible motifs. We sourced each SOX10 binding motif sequence from JASPAR^71^, SwissRegulon^72^, and HOCOMOCO^8^ and visualized each possible binding site at the *Hoxa6* locus (chr6-52206288-52208722). Visualization methods from Signac^73^ were used to visualize both genes and binding sites on this interval.

### Hybridization Chain Reaction with Immunolabeling

*In situ* hybridization chain reaction (HCR) was performed on isolated ileum samples from *Sox10^+/+^* and *Sox10^Dom/+^* fetal mice, as recommended by the manufacturer (Molecular Instruments) with minor modifications^74^. Briefly, fetal intestines were isolated and fixed overnight in 4% paraformaldehyde (PFA). Samples were subsequently dehydrated in methanol at −20°C. For HCR processing, samples were brought to room temperature and bleached in Dent’s bleach, washed and treated with Proteinase-K, then post-fixed in 4% PFA to ensure Proteinase-K inactivation. Samples were hybridized in hybridization buffer from the manufacturer (Molecular Instruments) overnight at 37°C and washed using recommended conditions. HCR probes targeting *Hoxa6* and *Elavl4* transcripts were designed using OligoMiner^75^ and screened for sequence similarity to gene family members in the mouse genome using NIH Basic Local Alignment Search Tool (BLAST) and UCSC BLAST-Like Alignment Tool (BLAT). Following HCR sample processing, whole-mount immunohistochemistry was performed after a 4% PFA post-fix step of samples using established methods for immunohistochemistry^76^.

| Type | Target/<br>Excitation | Variation | Probe<br>Count | Reference | Amount/250ul Hyb |
| --- | --- | --- | --- | --- | --- |
| Probe | *Hoxa6* | B1 | 17 | NM_010454.3 | 1 ul |
| Probe | *Elavl4* | B3 | 23 | NM_010488.4 | .8 ul |
| Amp | 647 | B1 | Na | Choi, et al.<br>2018 | 2.5ul |
| Amp | 488 | B3 | Na |  | 5 ul |

| Antibody | Target | RRID | Supplier | Dilution | Incubation |
| --- | --- | --- | --- | --- | --- |
| Primary | Hu C/D | AB_2813895 | Gift from<br>Vanda Lennon | 1:10,000 | Overnight,<br>4oC |
| Secondary | Donkey anti-Human,<br>Alexa 594 | AB_2340535 | Jackson<br>Immuno<br>709-495-149 | 1:1,500 | 4 hrs, RT |

### Image Capture

Confocal images of ileum tissue were acquired on a Zeiss LSM880 confocal with ZEN 2.3 (Black) software with the following parameters: green fluorophore imaged with 488 nm excitation and 493-598 nm emission (MBS 488); red fluorophore imaged with 561 nm excitation and 571-695 nm emission (MBS 458/561) and far red fluorophore imaged with 633 nm excitation and 638-755 nm emission (MBS 458/514/561/633). Tracks were acquired sequentially (frame). Objectives included Plan-Apochromat 63x/1.4 Oil; Plan-Apochromat 40x/1.3 Oil; and Plan-Apochromat 20x/0.8. For all images, the pinhole was set to 1 Airy Unit, Pixel Dwell Time was 4.1 used, Frame size was 1024×1024, and zoom =1. Laser and Gain settings were determined on the wild-type sample to optimize the dynamic range. All settings were then kept identical for each matched littermate sample. For each sample, a 20x image was first acquired by scanning the tissue starting at the distal region until HuC/D immunolabeling was located. One 20x image was acquired at this initial position. Next, 3 images at 40x were acquired in the same location and continuing proximally configured to avoid overlap of each image. Finally, one 40x or 63x image was acquired to show individual red/far red combination cells. Z range was set by starting at the area in the tissue where the *Elavl4* HCR signal began and focusing toward the coverslip to the slice where *Elavl4* HCR signal faded. Each Z-stack was approximately 8-12 um. Z-stacks were acquired with Nyquist sampling.

### Image Analysis

Images were imported into ImageJ and initially compressed into a Z-stack using the Z-stack function (average in-slices). Channels were separated using the Split Channel function to facilitate individual analysis of Hu C/D, *Hoxa6*, and *Elavl4* images and thresholding was independently conducted on each channel. Aiming to isolate true signal, HuC/D images were thresholded by 30-25%, *Elavl4* by 10-8%, and *Hoxa6* by 2.5-1.5% using the Otsu method. Regions of interest (ROI) were generated using Analyze Particle functions on the HuC/D channel, with a size limitation set at 50-infinity to focus analysis on regions occupied by ENPs. If single isolated HuC/D positive cells were present that were not captured by the 50-infinity size limit additional ROIs were set using the Virtual Wand Tool ImageJ plugin. Additional ROIs were set using the free hand tool within the sample image in space lacking HuC/D signal to serve as control regions for background noise subtraction. The percentage area was captured for all other channels using ROIs generated from the HuC/D channel. Images were normalized by subtracting the average of control ROIs, and HCR ratios were generated by dividing *Hoxa6* by *Elavl4* normalized percentage areas.

A step-wise reduction in threshold was applied to enable selective signal quantitation among dim HuC/D-positive cell populations. All cells were selected within the HuC/D channel using Image-J plugin Versatile Wand Tool and threshold reductions of 100%, 75%, 50%, and 25% were applied to limit ROIs to those areas with the 75% dimmest signal, the 50% dimmest signal, and the 25% least dim signal. Using the selected ROIs, percent areas were captured, and *Hoxa6* to *Elavl4* ratios were calculated by dividing *Hoxa6* by the *Elavl4* percent areas. For the 100% threshold, three control ROIs selected for minimal to no signal on the tissue to represent background were selected. These were then used to normalize the *Hoxa6* and *Elavl4* percent area values by taking the average of the percent area of the control ROIs per fluorescent channel and subtracting this value from each of the sample ROIs. To prevent generation of negative values, this process was not applied for the remaining signal threshold reductions (75%, 50%, and 25%). During generation of *Hoxa6* to *Elavl4* ratios, we excluded ROIs in which HuC/D antibody signal was present but *Elavl4* was not (ratio of *Hoxa6* to *Elavl4* in this case gives infinite values), no signal for either *Elavl4* or *Hoxa6* was present (gives NA values for dividing 0 by 0), and mean control area signal was greater than the sample area (gives negative signal values).

Three-dimensional renderings were created using IMARIS (Oxford Instruments, V10.1). Raw image files were initially converted to IMARIS files, preserving metadata through IMARIS image converter. In the 3D view, images were generated using the MIP (Max Image Projections) at the highest resolution quality. Videos were produced utilizing IMARIS Animation features. Images of the 3D view were generated with the Blend function and the highest rendering quality, emphasizing the overlap between channels in a snapshot.

### Statistical Analysis

All statistical analyses were conducted following the standard methods of each respective R package. One-way Analysis of Variance (ANOVA) was employed using the R avo function to evaluate the statistical significance of cell count abundance across genotypes within each cluster. Cell counts were extracted from Seurat objects for each *Sox10^+/+^* and *Sox10^Dom/+^* biological replicate and grouped into clusters for this analysis. *P*-values below 0.05 were considered statistically significant.

Since scRNA-seq expression profiles show that *Hoxa6* is only expressed in some and not all clusters, normalized HCR ratios of *Hoxa6*:*Elavl4* percent area for each of the four signal percentages were filtered for positive values. These *Hoxa6*:*Elavl4* percent area ratios were heavily skewed towards low values, so the data were log10-transformed to bring the distributions closer to normal/Gaussian. The Shapiro-Wilk tests for normal distributions revealed that these data distributions were still non-normal. Therefore, a Wilcoxon non-paired test (non-parametric t-test equivalent for independent samples) was applied to test differences between HCR *Hoxa6*:*Elavl4* ratios of WT and *Sox10^Dom^* fetal intestine tissues for each signal threshold reduction (Figure 8E). *P-*values below 0.05 were considered statistically significant.

## Abbreviations

CNS: Central nervous System
dpc: Days Post-Coitus.
ENS: Enteric nervous system
ENCP: Enteric Neural Crest-Derived Progenitors
GI: Gastrointestinal
GO: Gene Ontology
GRN: Gene Regulatory Networks
HSCR: Hirschsprung’s disease
HCR: Hybridization Chain Reaction
NC: Neuronal Cell
PC: Progenitor Cell
PFA: Paraformaldehyde
ROI: Regions of Interest
SCENIC: Single-Cell Regulatory Network Inference and Clustering
scRNA-seq: single-cell RNA sequencing
UMAP: Uniform Manifold Approximation and Projection analysis.

## Transcript Profiling

**single cell** RNA-seq data from this study have been deposited into the National Center for Biotechnology Information Gene Expression Omnibus (NCBI, GEO) under accession number: GSE262898.

## Data access

R Code for data processing and R Objects are provided at Open Science Framework.

R Code, ImageJ Output, and Tables: https://osf.io/bkfmg/?view_only=8fd67a70615348d89d6618fabbeec033

Seurat object containing all cells: https://osf.io/h8s4b/?view_only=1881bb55f0ab4c318c627f9397f562d5

Seurat object containing only Neuronal Cells: https://osf.io/58cr4/?view_only=2f5d9d62673c4276849861c34e165675

Seurat object containing only Progenitor cells: https://osf.io/br3ac/?view_only=1f757e7348cd4ed18fe0811b81ccfb2d

CellRanger Output: https://osf.io/hsbf2/?view_only=b488dfacb17848019196215157a4bc9d

## Author Contributions

EMS^2^ and JAA designed the studies. All authors were critically involved in data collection. JAA and JTB performed data and statistical analyses. JAA, JTB and EMS^2^ interpreted data. JAA and EMS^2^ drafted the manuscript. EMS^2^ and JTB edited the manuscript. JTB generated shared data resource links. EMS^2^ obtained funding. EMS^2^ supervised the study. All authors revised and approved the manuscript.

## Synopsis

Single cell transcriptomics in *Sox10^Dom^*Hirschsprung mice identifies defects in specification of enteric neurons during development. Analysis of gene regulatory networks reveals *Hox* genes, prominently *Hoxa6*, as regulators of enteric neuron diversity. Knowledge of ENS gene regulatory networks facilitates directed differentiation of enteric neurons for treatment of bowel motility disorders.

## Acknowledgments

The authors thank the support staff of the Cell Imaging Shared Resource (CISR) Core at Vanderbilt for advice and assistance in confocal imaging and image analysis with IMARIS. Confocal microscopy was performed in the Cell Imaging Shared Resource (CISR) Core at Vanderbilt supported National Institutes of Health Grants (CA68485, DK20593, DK58404, DK59637, EY08126, and S10 OD021630). Imaging was performed with partial support from a Core Scholarship funded by P30DK058404 via the Vanderbilt University Medical Center’s Digestive Disease Research Center. HuC/D antibody was a kind gift from Vanda Lennon (Mayo Clinic). We are grateful to Dr. Rosa Uribe for recommendations on immunolabeling after HCR.

